# Females do not avoid but prefer superparasitism in the parasitoid wasp *Melittobia australica*

**DOI:** 10.64898/2026.10.13.758990

**Authors:** Iris Veenis, Bart A. Pannebakker, Jun Abe

**Affiliations:** Laboratory of Genetics, Wageningen University, Wageningen, The Netherlands; Faculty of Sciences, Kanagawa University, Kanagawa, Japan

**Keywords:** foraging behaviour, host diapausing status, ideal free distribution, local resource enhancement, *Melittobia australica*, parasitism efficiency, parasitoid wasp, sex allocation, superparasitism

## Abstract

To reduce competition over limited resources, organisms often avoid resources already used by others. In parasitoid insects, females are generally expected to avoid parasitizing hosts already parasitized by other females, called superparasitism, and this prediction has been broadly supported across numerous species. However, in the parasitoid wasp *Melittobia australica*, multiple females were observed ovipositing on the same host despite the presence of nearby unparasitized hosts in the field. In this study, we first analysed field data using randomization tests, and found that females strongly aggregated on particular hosts rather than distributing randomly across hosts. We then conducted laboratory experiments to examine (1) whether this aggregation reflects an active preference for superparasitism and (2) whether such behaviour could improve parasitism efficiency. A choice experiment, in which females were provided with a parasitized and an unparasitized host, showed that females preferred superparasitism when hosts were non-diapausing. This result indicates that females of *M. australica* not only engage in superparasitism, but actively prefer parasitized hosts. Another experiment, in which varying numbers of females were allowed to oviposit on a host, found that offspring number per female decreased with increasing female number on diapausing hosts, whereas it remained constant irrespective of female number on non-diapausing hosts. This host-dependent result suggests that multiple females can more effectively paralyze vigorous non-diapausing hosts, thereby offsetting the negative effects of offspring competition. As *Melittobia* females are likely to attack non-diapausing hosts in most cases in the field, including those in the present study, this benefit of parasitism by multiple females may help explain their preference for superparasitism, in contrast to many other parasitoid species.

## Introduction

In nature, organisms often compete for limited resources. To reduce competition and maximize their fitness, individuals adjust their distribution and foraging behaviour in response to resource availability. The ideal free distribution theory predicts that individuals spread themselves according to the abundance of resources and competitors to minimize competition, and this prediction has been broadly supported across numerous species (Fretwell & Lucas 1970; Kacelnik et al. 1992; Křivan et al. 2008). This applies broadly to organisms searching for spatially distributed resources, including food, breeding sites, and other resources essential for reproduction (Matsumura et al., 2010).

Parasitoids are insects that lay their eggs on or inside arthropod hosts, usually insects (Godfray, 1994). Since parasitoid offspring rely entirely on their host for all resources required for their development, host selection by their mothers (hereafter called foundresses) directly determines the fate of their development. When parasitoid foundresses oviposit in or on hosts that have been previously parasitized by other conspecific foundresses, called superparasitism, competition for host resources occurs among offspring of these different foundresses. In gregarious parasitoids, scramble competition reduces the number of emerging offspring, and in solitary parasitoids, contest competition typically results in the emergence of only a single individual (Visser 1996; Tuda & Iwasa 1998). To prevent the disadvantage in such competition, parasitoid foundresses are predicted to avoid superparasitism, and empirical studies have shown that females of most parasitoid species do so (Godfray 1994; Tregenza et al. 1996; Fauvergue et al. 2006; Dorn & Beckage, 2007; Ode et al. 2022; Fellows et al 2023).

Although superparasitism has negative consequences and is generally avoided, it nevertheless occurs in nature. Parasitoid foundresses may adopt superparasitism as a suboptimal choice when hosts are scarce in the habitat and unparasitized hosts are not available (van Alphen & Visser 1990). Moreover, there are conditions under which superparasitism can be adaptive. For example, in solitary endoparasitoids, self-superparasitism or depositing multiple-egg clutches in one host can increase the probability that an offspring successfully completes development within a host by suppressing host immune defence (Rosenheim and Hongkham 1996; Khafagi and Hegazi 2008; Liu et al. 2018). In a facultatively gregarious parasitoid wasp *Coccus hesperidum*, laying a larger number of eggs in a host decreases the encapsulation rates of the eggs and increases offspring survival (Kapranas et al. 2012). In addition, effective host suppression by multiple foundresses has been reported in gregarious ectoparasitoid wasps of the genus *Sclerodermus*, in which superparasitism can improve the success rate of parasitism and increase offspring production (Tang et al. 2014; Abdi et al. 2020a, b; Liu et al. 2021).

*Melittobia australica* is an ectoparasitoid wasp attacking mainly the prepupae of solitary bees and wasps, in which foundresses lay eggs on the surface of the hosts body and the offspring feed on the hosts hemolymph from outside (Matthews et al., 2009; Dahms, 1984). *Melittobia australica* is a gregarious parasitoid that potentially lays several hundred eggs under favourable laboratory conditions, and around one hundred eggs even under unstable field conditions (Abe et al. 2003; 2021). Hosts of *Melittobia* are clumped within host nests, and *Melittobia* foundresses that arrive at a host nest choose a host on which to lay eggs. Interestingly, foundresses of *M. australica* are sometimes found to aggregate on a single host, despite the presence of numerous unparasitized hosts within host nests (Abe et al. 2021).

In this study, our aim was to investigate whether *M. australica* foundresses prefer to oviposit on hosts with other foundresses, and how superparasitism influences parasitism efficiency. First, using field data, we examined whether multiple foundresses tend to lay eggs on the same host using a randomization approach. However, this analysis could not distinguish whether females actively prefer aggregation or simply aggregate incidentally on particular hosts. In the first experiment, we therefore conducted a choice test to examine whether foundresses prefer parasitized over unparasitized hosts when both were available. Using microsatellite DNA markers, we identified which hosts the focal foundresses laid eggs on and how many offspring they produced. In the second experiment, we examined how the number of females ovipositing on a host affects parasitism efficiency by comparing individual offspring production. In both experiments, we used non-diapausing hosts that are active and may resist parasitism, and diapausing hosts that are not active, to examine how host status influences foundress behaviour and parasitism efficiency.

## Materials and Methods

### Parasitoid wasp and host species

In *Melittobia*, mating occurs among individuals that developed on the same host, and only females disperse from the natal host to search for new hosts. Females exhibit two distinct wing morphs: a minority of short-winged morphs and a majority of long-winged morphs (Dahms 1984; Matthews et al. 2009). In the present experiments, we only used long-wing females for foundresses. Once *Melittobia* foundresses find suitable hosts, they inject a venom that halts development and paralyzes the host before laying eggs (Deyrup et al., 2006). In the meanwhile, they consume a portion of the host hemolymph to mature their eggs (synovigeny), after which they begin oviposition on the host (Mathews et al. 2009). In the experiments described below, we used two strains of *M. australica*: the IA13 strain, collected in Ishigaki, Okinawa, Japan, in 2013, and the L strain, collected in Otsu, Shiga, Japan, in 2000 (Abe et al. 2005). These strains have been maintained in the laboratory since collection, and females were randomly drawn from stock cultures for use in the experiments. The two strains can be distinguished by genotyping individuals using microsatellite markers (Abe and Pannebakker 2017).

*Melittobia* species are generalists and parasitize a wide range of solitary wasps and bees that nest above ground (Matthews et al. 2009). Their hosts can be parasitized from the larval to pupal stages, but they preferentially parasitize prepupae. In the present experiments, we used prepupae of the giant leaf-cutter bee, *Megachile sculpturalis* (Hymenoptera: Megachilidae), the most common host of *Melittobia* species in field surveys using bamboo traps (Abe et al. 2021). The prepupa of *M. sculpturalis* is typically much larger than an individual of *M. australica*, and several thousand offspring can potentially emerge from a single host prepupa when multiple females oviposit on it under laboratory conditions (Abe et al. 2003). The hosts used were either diapausing or non-diapausing prepupae. Diapausing prepupae were collected in December 2023 in Hiratsuka, Kanagawa, Japan, and stored at approximately 4°C in a refrigerator until the experiments. Non-diapausing prepupae were collected in November 2024 in the same area and used immediately in the experiments. Before the experiments, host prepupae were weighed using a Libror AEX-200G scale (Shimadzu, Japan). All the experiments below were conducted at 25°C and with 0h light/24h dark cycle.

### Distribution of foundresses in the field

To determine whether *M. australica* foundresses lay eggs on the same host, we analysed field data on the distribution of foundresses across hosts. The data of the number of foundresses per host was determined by sampling foundresses on hosts using bamboo traps in the field, and genotyping offspring using microsatellite markers (Abe et al. 2021). Foundresses of *M. australica* can adopt one of two dispersal behaviours: dispersers, which fly to other patches to find new hosts, and non-dispersers, which remain and search for new hosts within their natal patches (Abe et al. 2021). Foundress dispersal behaviours are not necessarily consistent with female wing morphs, although the short-winged morph is always of the non-disperser type. Here, we analysed only dispersing foundresses and excluded non-dispersers, because our aim was to examine host selection by foundresses dispersing independently. Non-dispersing foundresses tend to arrive at hosts located near their natal hosts within host nests, and thus their distribution among hosts is likely to reflect natal location more strongly than host choice.

To assess whether foundresses are significantly aggregated across hosts, we performed randomization tests, in which the observed data were compared with the distribution generated from repeated simulations that randomly assigned foundresses to hosts. The P value was calculated as the proportion of simulation iterations that produced values as extreme as or more extreme than the observed value. We used two datasets from Abe et al. (2021) for these tests. First, we used data on all hosts from all patches (Table 1A), treating all hosts independently of patch identity. The observed superparasitism rate among parasitized hosts was compared with a null distribution generated from 2,000 simulation iterations. In each iteration, 31 foundresses (matching the observed foundress numbers) were randomly and independently assigned to one of 4,871 hosts, and the superparasitism rate was calculated. In the second randomization, we used data from seven patches in which more than one female arrived (Table 1B). This analysis focused on the foraging behaviour of foundresses that arrived at patches where other foundresses were also present. The observed mean number of foundresses per host among the seven patches was compared with the corresponding null distribution (2,000 simulation iterations). In each iteration, the same number of foundresses as observed were randomly assigned to the same number of hosts within each patch, and mean numbers of foundresses were calculated.

**Table 1.** Distribution of the number of hosts parasitized by different numbers of *Melittobia australica* foundresses in the field.

|  | Foundress number on a host |  |  |  |  |  |
| --- | --- | --- | --- | --- | --- | --- |
|  | 0 | 1 | 2 | 3 | 4 | 5 |
| A. All patches <sup>1</sup> | 4840 | 23 | 6 | 1 | 0 | 1 |
| B. Patches with multiple foundresses (7 patches) <sup>1</sup> |  |  |  |  |  |  |
|  | 61 | 1 | 2 | 0 | 0 | 0 |
|  | 51 | 0 | 0 | 0 | 0 | 1 |
|  | 35 | 0 | 0 | 1 | 0 | 0 |
|  | 18 | 0 | 1 | 0 | 0 | 0 |
|  | 30 | 0 | 1 | 0 | 0 | 0 |
|  | 42 | 0 | 1 | 0 | 0 | 0 |
|  | 60 | 0 | 1 | 0 | 0 | 0 |

<sup>1</sup> Patches represent bamboo traps used to provide nesting sites for host species. Data from Abe et al. (2021).

### Experiment 1: preference for unparasitized versus parasitized hosts

We tested whether *M. australica* foundresses prefer parasitized hosts over unparasitized hosts for oviposition. We allowed a foundress, called “focal foundress”, to choose between a host parasitized by another foundress, called “treatment foundress”, and an unparasitized host. To determine which hosts the focal foundress laid eggs on and how many offspring each female produced, we used different strains for the focal and treatment females, and genotyped their offspring using microsatellite markers. Both the IA13 strain and the L strain were used for both focal and treatment foundresses.

A treatment foundress was introduced into a plastic case (86 mm in diameter, 30 mm in height) containing a host prepupa in a plastic cup (24 mm in diameter, 10 mm in height). The foundress was checked daily for oviposition behaviour under minimal light conditions. When her abdomen began to enlarge, she was considered to have started oviposition (Matthews et al. 2009), and was allowed to lay eggs for an additional three days. After this period, a cup with an unparasitized host was added to the case and the focal foundress was introduced, allowing the focal foundress to choose between the parasitized and unparasitized hosts. The parasitized and unparasitized hosts in the cups were placed approximately 20 mm apart in the plastic case, and the two hosts were either both diapausing or both non-diapausing.

After a further five days, both foundresses were removed, and one of the hosts was transferred to another plastic case with the cup. The hosts were maintained until the offspring emerged, and the emerged offspring were counted and preserved in 99.5% ethanol. If offspring developed from both hosts, we genotyped 8 offspring randomly selected from each host and determined whether offspring from each host were produced by the treatment or focal foundresses of different strains. If offspring developed from only one host, we genotyped 16 offspring from the host and estimated the number of offspring produced by each foundress based on the proportion of the two strains among the 16 individuals. Replicates in which focal foundresses laid no eggs on either the parasitized or unparasitized host were excluded from the analysis (two and two for the diapausing and non-diapausing host treatments, respectively). Consequently, eight and nine replicates remained for the diapausing and non-diapausing host treatments, respectively.

### Molecular analysis

DNA was extracted from the head and thorax of adult females to avoid contamination from sperm stored in the spermatheca. DNA extraction was performed using the boiling method, as described elsewhere (Abe et al. 2009). To distinguish individuals of the L and IA13 strains, PCR amplification was performed using one of four primer sets (Mau15, Mau17, Mau19, or Mau61) for each individual (Abe and Pannebakker 2017). PCR was conducted using the M13-tailed primer technique (Schuelke 2000) in a 5 μL reaction mixture containing 0.5 μL primer stock (0.05 μM forward and 0.2 μM reverse primers), 0.5 μL M13 primer (10 μM) labelled with a fluorescent dye (6-FAM, VIC, NED, or PET), 0.5 μL Type-it MasterMix (Qiagen), and 1 μL genomic DNA. Amplification was carried out using an ABI 2720 thermal cycler (Life Technologies, Carlsbad, CA, USA) under thermal cycling conditions described elsewhere (Abe and Pannebakker 2017). The amplified fragments were analysed using Peak Scanner software version 1.0 (Life Technologies, Carlsbad, CA, USA) following electrophoresis on an ABI 3130 capillary sequencer (Life Technologies, Carlsbad, CA, USA). The four primer sets were labelled with different fluorescent dyes and amplified in separate PCR reactions, but PCR products generated using different primer sets were run simultaneously by electrophoresis, reducing the number of electrophoresis runs.

### Experiment 2: the effect of foundress number on offspring production

To determine how superparasitism influences the efficiency of offspring production, we varied the number of foundresses (one, four, or eight) and allowed them to oviposit on a host, measuring the number of emerged offspring. To examine how the effect depends on host diapause status, we used both diapausing and non-diapausing hosts. Therefore, this experiment employed a 3 × 2 factorial design incorporating female number and host status as factors. Females from either the L or IA13 strain were used for foundresses in each replicate, and three replicates were conducted for each strain across all six treatment combinations.

Either one, four, or eight foundresses were introduced into a plastic case with a host prepupa and were allowed to lay eggs on the host. After five days, the number of live foundresses were counted, and all the foundresses were removed. The host was then maintained until offspring emergence, after which the emerged offspring were sexed and counted. For the treatments with four and eight females, the average number of offspring produced per female was calculated by dividing the total number of offspring by the number of females introduced. The sex ratio for each replicate was calculated as the proportion of males among the total number of offspring per host in that replicate.

### Statistics

All analyses were conducted using R version 4.4.2 (The R Foundation for Statistical Computing, Vienna, Austria). Data on host choice probability and survival probability were analysed using generalized linear models (GLMs) fitted with binomial distributions. To account for overdispersion, data on sex ratio and offspring number were analysed using generalized linear mixed models (GLMMs) fitted with binomial and Poisson error distributions, respectively, with replicate identity included as a random effect. Full models initially included all fixed effects and interaction terms of interest, and were subsequently simplified by backward stepwise elimination. Terms were removed starting with higher-order ones, and model comparisons were conducted using likelihood ratio tests. Non-significant terms (α > 0.05) were sequentially removed from the least significant term to obtain the minimal adequate model.

## Results

### Distribution of foundresses in the field

Across all patches, the average superparasitism rate observed in the field (0.258) was significantly higher than that expected under the null distribution generated by the randomization test (mean ± SD = 0.0032 ± 0.0101; P < 0.001). In patches with multiple foundresses, the observed mean number of foundresses on a host (2.81) was also significantly higher than that expected under the null distribution (0.0575 ± 0.0744; P < 0.001). These results suggest that *M. australica* foundresses do not distribute randomly across hosts, but instead aggregate on particular hosts.

### Experiment 1: preference for unparasitized versus parasitized hosts

Microsatellite data showed that treatment foundresses produced all their offspring on the initially provided hosts, and focal foundresses produced all their offspring on either the parasitized or the newly provided unparasitized host. No mixed clutches across the two hosts were observed. Diapausing host prepupae showed little movement, whereas non-diapausing prepupae wriggled actively, specifically in contact with parasitoid females. However, once foundresses progressed in oviposition, movement decreased even in non-diapausing prepupae. The probability that focal foundresses chose parasitized versus unparasitized hosts was not influenced by weight differences between the two hosts (GLM: χ^2^_1_ = 0.094, P = 0.759), although the strain of focal females and host diapause status showed marginal effects (strain: χ^2^_1_ = 3.18, P = 0.074; host status: χ^2^_1_ = 3.23, P = 0.072). Focal foundresses provided with diapausing hosts did not show significant preference between parasitized and unparasitized hosts (binomial test: P = 1.00), those provided with non-diapausing hosts significantly preferred immobile parasitized hosts over unparasitized hosts to lay eggs (Figure 1; binomial test: P = 0.039).

**Figure 1.**
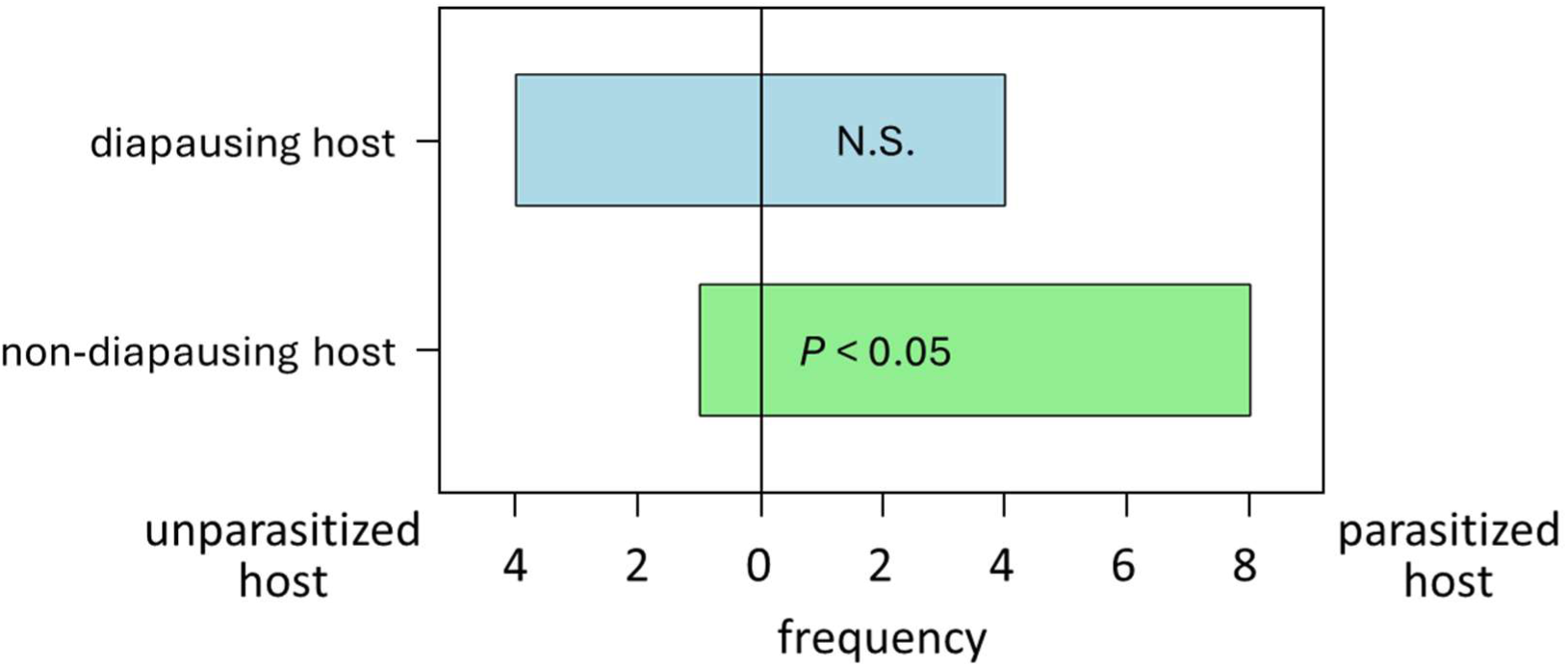
Frequency with which focal foundresses chose unparasitized or parasitized hosts. The two hosts provided in each replicate were either both diapausing or both non-diapausing. *P* values shown on the bars indicate significant differences in the probability of choosing between unparasitized and parasitized hosts (binomial test).

Treatment foundresses produced a significantly larger number of offspring when they laid eggs on diapausing hosts than on non-diapausing hosts (Figure 2a; GLMM: χ^2^_1_ = 9.75, P < 0.01), and foundresses from the L strain produced more offspring than those from the IA13 strain (χ^2^_1_ = 5.01, P = 0.025). The number of offspring was not significantly influenced by whether a focal foundress subsequently joined in oviposition on the same host or not (Figure 2a; interaction between host parasitism status and host diapause status: χ^2^_1_ = 0.971, P = 0.324; host parasitism status: χ^2^_1_ = 0.160, P = 0.689), and host weight showed only a marginal effect (χ^2^_1_ = 3.08, P = 0.079). Focal foundresses also produced a significantly larger number of offspring when they laid eggs on diapausing hosts than on non-diapausing hosts (Figure 2b; GLMM: χ^2^_1_ =6.63, P = 0.01). However, offspring number was not influenced by whether the foundresses chose previously parasitized versus unparasitized hosts (Figure 2b; interaction between host parasitism status and diapause status: χ^2^_1_ =0.103, P = 0.748; host parasitism status: χ^2^_1_ = 0.103, P = 0.748), nor was it influenced by other factors (host weight difference: χ^2^_1_ = 2.18, P = 0.140; focal foundress strain: χ^2^_1_ = 2.63, P = 0.105).

**Figure 2.**
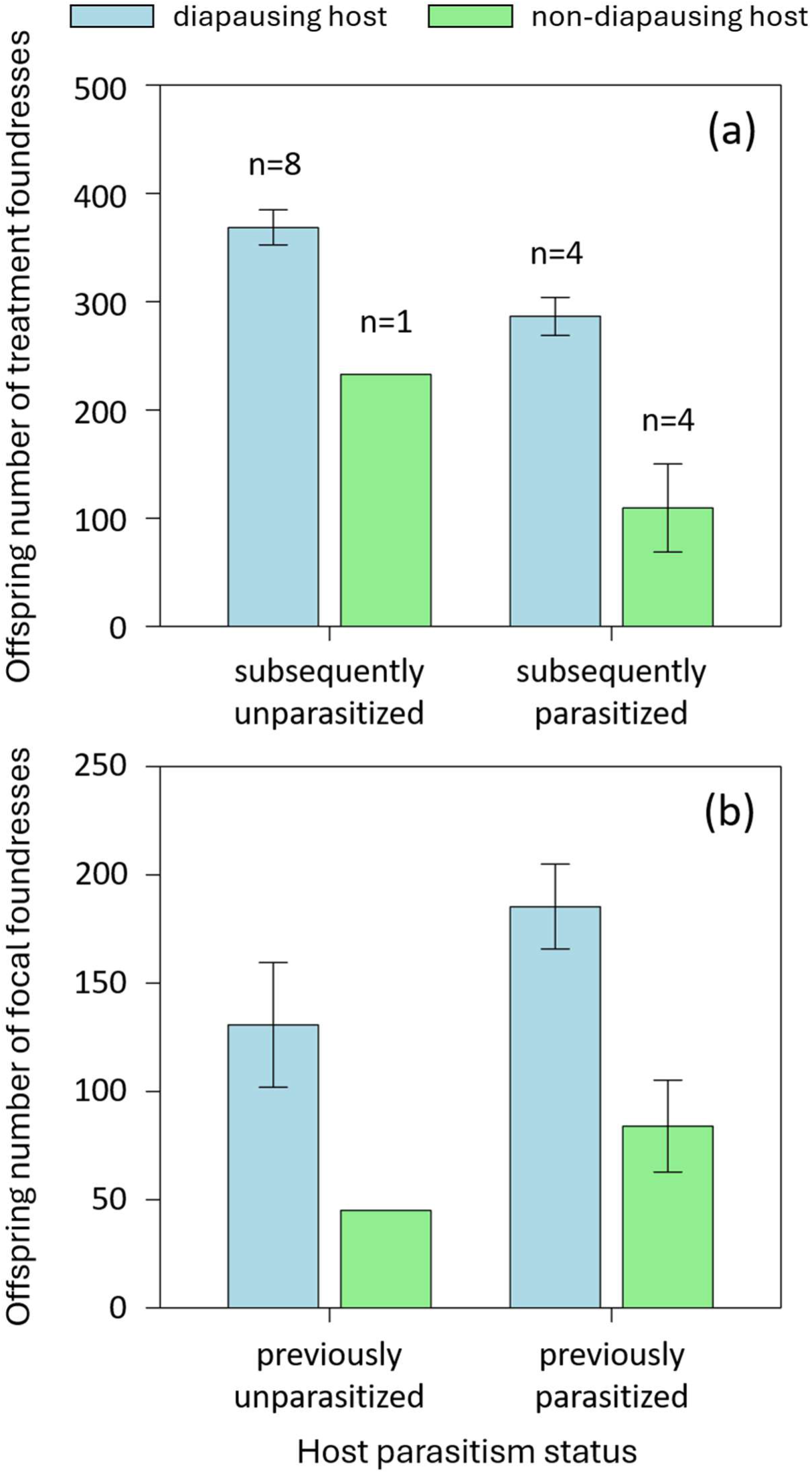
Average number of offspring produced by a treatment foundress depending on whether a focal foundress subsequently joined in oviposition on the same host or not (a), and that produced by a focal foundress depending on whether a treatment foundress had already been ovipositing on the same host or not (b). The two hosts used in each replicate were either both diapausing or both non-diapausing. Error bars indicate standard errors. The number of replicates above bars in (a) are the same for (b).

### Experiment 2: the effect of foundress number on offspring production

A proportion of foundresses (0.032 foundresses in replicates overall) had died by 5 days after the introduction. However, the survival probability of the foundresses was not affected by any of the tested effects (GLM: interaction between foundress number and host diapause status: χ^2^_1_ = 0.346, P = 0.557; foundress number: χ^2^_1_ = 0.149, P = 0.700; host diapause status: χ^2^_1_ = 0.817, P = 0.366; host weight: χ^2^_1_ = 2.36, P = 0.124; foundress strain: χ^2^_1_ = 0.328, P = 0.567).

The average number of offspring produced per foundress was significantly influenced by the interaction between foundress number and host diapause status (Figure 3a; GLMM: χ^2^_1_ = 5.59, P = 0.018), while other factors were non-significant or marginal (host weight: χ^2^_1_ = 0.006, P = 0.938; female strain: χ^2^_1_ = 3.20, P = 0.076). When the two host diapause status were analysed separately, offspring production per foundress on diapausing hosts significantly decreased with increasing foundress number (χ^2^_1_ = 7.39, P < 0.01). However, offspring production per foundress on non-diapausing hosts did not change significantly despite increasing foundress number (χ^2^_1_ = 0.566, P = 0.452). The sex ratio of the offspring (proportion of males) increased slightly but significantly with foundress number (Figure 3b; GLMM: χ^2^_1_ = 16.7, P < 0.001). The remaining effects had no significant or only marginal effects on offspring sex ratio (interaction between foundress number and host diapause status: χ^2^_1_ = 0.001, P = 0.975; host diapause status: χ^2^_1_ = 0.091, P = 0.763; host weight: χ^2^_1_ = 0.149, P = 0.70; foundress strain: χ^2^_1_ = 3.35, P = 0.067).

**Figure 3.**
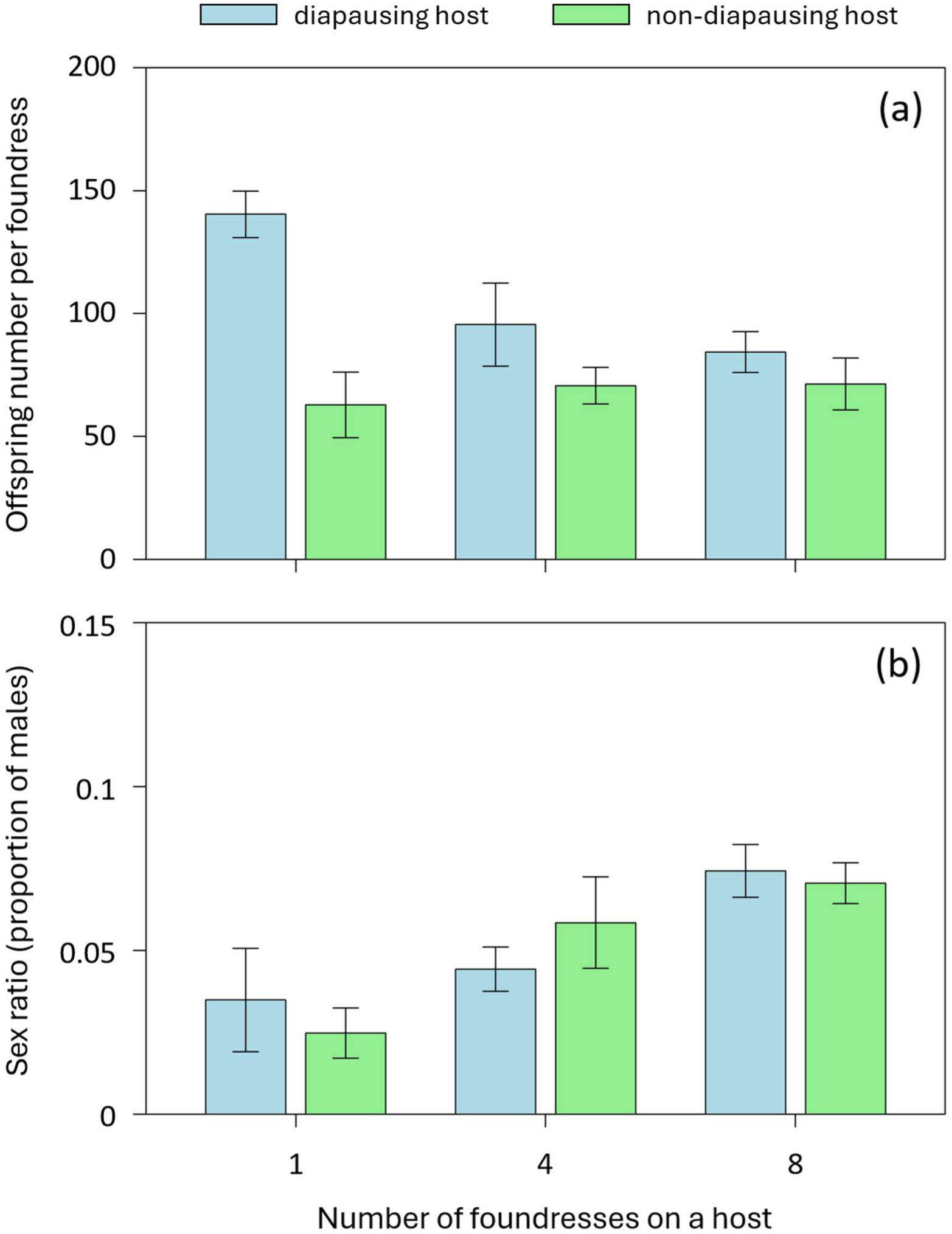
Average number of offspring per foundress (a) and offspring sex ratio (proportion of males; b) when allowed to oviposit on a host alone or in groups of four or eight females. Hosts were either diapausing or non-diapausing. Error bars indicate standard errors. The number of replicates for all treatments was six.

## Discussion

The present study investigated whether foundresses of *M. australica* prefer to oviposit together on the same host and, if so, why they engage in such superparasitism. Analyses of the field data indicated that foundresses tended to aggregate on particular hosts. This tendency could also result from preferences for particular hosts (e.g. due to host quality) or limited availability of hosts that foundresses could encounter. However, our first experiment showed that foundresses preferred hosts already parasitized by another foundress when hosts were non-diapausing. In addition, our second experiment showed that offspring production per foundress did not decrease even with increasing foundress number on non-diapausing hosts. Broods in the field samples analysed were likely initiated before November (Abe et al. 2021), when hosts were still non-diapausing, and most *Melittobia* females are likely to attack non-diapausing hosts in the field. Therefore, the observed tendency toward superparasitism in *M. australica* may be associated with the absence of reduced offspring production on non-diapausing hosts.

In stark contrast to the theory and behaviour reported for numerous other parasitoid wasp species (Godfray 1994; Ode et al. 2022; Fellows et al 2023), the present study showed that *M. australica* foundresses prefer superparasitism depending on host conditions. As shown by the second experiment, offspring number per foundress decreased with increasing foundress number on a diapausing host, which is typically observed in many gregarious parasitoid species and as also shown in *Melittobia* (Godfray 1994; Abe et al. 2003). This indicates that competition occurs among offspring for host nutrients. In contrast, no decrease in offspring production was observed on non-diapausing hosts, suggesting that communal oviposition by multiple foundresses outweighs the negative effects of offspring competition. Because non-diapausing hosts move frequently, they may be difficult and risky for foundresses to parasitize. Oviposition by multiple foundresses could therefore facilitate host paralysis and immune suppression. Specifically, when foundresses arrive at hosts sequentially, those that choose hosts already parasitized by others may gain access to hosts that have already been suppressed.

The first experiment showed that foundresses prefer parasitized hosts on non-diapausing hosts, and this behaviour could arise from two different mechanisms. Foundresses may actively prefer to oviposit together because this facilitates host suppression, or they may simply prefer already parasitized hosts because parasitized hosts are less mobile. In the field, *Melittobia* foundresses are reported to reach suitable hosts in two different ways: they can invade host nests after hosts reach an appropriate developmental stage, or they can remain inside nests and wait for hosts to mature (Dahms 1984; Matthews et al. 2009). These different foraging patterns could influence how multiple foundresses encounter and oviposit on the same host under natural conditions. Future studies involving detailed observations of foraging behaviour in the laboratory, as well as investigations of how multiple foundresses locate and approach suitable hosts in the field, would help disentangle the two possible explanations for females’ preference for superparasitism.

Similar cooperative host suppression has been documented in bethylid parasitoid wasps, *Sclerodermus* species. Multiple foundresses cooperate to paralyze large and defensive hosts (wood-boring coleopteran pupae), resulting in higher reproductive success and sometimes higher offspring production per foundress (Tang et al. 2014; Abdi et al. 2020a, b; Liu et al. 2021). Even after host suppression, foundresses also cooperatively care for offspring developing on the host (Hu et al. 2012), in contrast to *Melittobia* species, in which no parental care or other interactions with juveniles have been documented. In addition, laboratory experiment showed that multiple foundresses of *Sclerodermus* tend to aggregate on particular hosts (Malabusini et al. 2025), although they frequently move between hosts in contrast to *Melittobia* foundresses.

In *Sclerodermus*, whereas foundresses initially accept newly arriving other foundresses on the same host, they later exhibit aggressive behaviour toward other foundresses, sometimes resulting in the death of the opponents (Guo et al. 2023). In *Melittobia*, aggressive interactions are also observed between foundresses (Matthews and Deyrup 2007; Abe et al. 2021), although they appear to be rare, and were not observed in the present study. While multiple foundresses commenced oviposition without aggression, after eight days some foundresses were observed to bite and cut off the antennae and legs of rival foundresses, forcing them to leave the host (Abe et al. 2021). During the early stage of oviposition, when host resources are abundant, foundresses may cooperatively paralyze hosts without aggression. However, as resources become limited, foundresses may exhibit aggressive behaviour to exclude rival foundresses from the host.

Although communal oviposition in *M. australica* is likely to counteract the negative effects of competition among developing offspring on non-diapausing hosts, reproductive success per foundress did not improve through superparasitism. Why do *M. australica* foundresses still prefer superparasitism? First, the condition of foraging foundresses and environmental conditions may be harsher in the field than under stable laboratory conditions. In the field, failed broods and broods producing only a small number of offspring are frequently observed (Abe et al. 2021). Under such conditions, parasitism by multiple foundresses may be more effective. Second, the effectiveness of multiple foundresses may depend on the specific conditions. In *Sclerodermus*, parasitism efficiency varies with kinship between foundresses and host characteristics (Tang et al. 2014; Abdi et al. 2020a, b; Liu et al. 2021). Moreover, studies on the same species, *S. guani* (= *harmandi*), reported contrasting results regarding relationship between offspring production per foundress and the number of foundresses (Tang et al. 2014; Liu et al. 2021). In *Melittobia*, which are generalists, variation in host size and species may also influence the effectiveness. While small hosts may lead to greater competition among multiple foundresses, large hosts are likely to be more effectively suppressed by multiple foundresses. In addition, multiple foundresses may have an advantage against the bodyguarding mite species *Ensliniella parasitica*, which lives symbiotically with the host wasp *Allodynerus delphinalis* and attacks *Melittobia* foundresses to defend the host (Okabe and Makino 2008). Third, parasitism by multiple foundresses may provide benefits other than improving parasitism efficiency. For example, as multiple females in *Melittobia* are reported to cooperatively open one tunnel (Deyrup et al. 2005), this behaviour may help foundresses access hosts within nests

Sex ratios observed in *Melittobia* species remain puzzling. The extremely female-biased sex ratio increases little with the number of foundresses (Abe et al. 2003; 2014), as also shown in the present study, contradicting the prediction of local mate competition (LMC) theory (Hamilton 1967). Previous work partly solved this by showing that dispersing foundresses increase the proportion of sons, whereas non-dispersing foundresses do not (Abe et al. 2021). Although a model incorporating female dispersal status qualitatively explained the observed pattern, observed sex ratios remained more female-biased than predicted (Abe et al. 2021). Communal oviposition has been suggested as one possible explanation because it may promote cooperation among related females (local resource enhancement, LRE; Abe et al. 2021; Iritani et al. 2021). However, the present study suggests that this effect may be limited, as communal oviposition did not substantially improve reproductive success per foundress. In addition, theory does not predict that such behaviour would strongly bias sex ratios (Iritani et al. 2021). Similarly, *Sclerodermus* wasps show more female-biased sex ratios than predicted by LMC theory (Tang et al. 2014; Malabusini et al. 2026), and LRE has been proposed as an explanation for this pattern (Tang et al. 2014; Kapranas et al. 2016). However, its effect may be limited (Iritani et al. 2021), and alternative factors have subsequently been proposed (Lehtonen et al. 2023). Future studies integrating superparasitism, female interactions, kinship, foraging behaviour, and sex allocation will provide further insights into the unusual reproductive biology of these parasitoid wasps.

## Data Availability

Data for this study are available within the Supplementary Material.

## Acknowledgement

We would like to thank Hiroaki Yanagisawa for his preliminary work, Momoko Ichinokawa for her input on the analysis of the field data and Kazuo Takahashi (Kanagawa University) for granting access to his laboratory. This work was funded by the Uyttenboogaart-Eliasen Stichting Research Internship grant, Stichting Jo Kolk Studiefonds MSc. Internship Research grant, Stichting Fonds Dr. Christine Buisman MSc. Internship grant, Erasmus+ Traineeship grant and Stichting de Korinthiërs grant to I.V. and by Japan Society for the Promotion of Science KAKENHI (grant no. 21K06353, 21KK0267) to J.A..

